# The small GTPase Rab5 inhibits actin polymerization mediated by the *Legionella pneumophila* effector VipA

**DOI:** 10.1101/2025.02.18.638817

**Authors:** Joana Saraiva, Joana N. Bugalhão, Teresa Carvalho, Zach Hensel, Alvaro Crevenna, L. Jaime Mota, Irina S. Franco

## Abstract

*Legionella pneumophila* is a facultative intracellular Gram-negative bacterium that causes Legionnaires’ disease, one of the most severe manifestations of atypical pneumonia. *L. pneumophila* naturally thrives within protozoan hosts in aquatic environments, which serve as reservoirs for bacterial replication and transmission. Inhalation of contaminated aerosols delivers *L. pneumophila* to the lungs where it is able to infect human alveolar macrophages and multiply intracellularly inside a membrane-bound compartment, the *Legionella*-containing vacuole (LCV). The main virulence factor of the bacteria is the Icm/Dot type IVb secretion system, responsible for the translocation of over 300 bacterial effector proteins into host cells. These effectors allow the remodelation of the LCV into a replication-competent compartment that seggregates from the phagolysosomal pathway.

After being translocated into host cells, *L. pneumophila* effector VipA associates with early endosomes and F-actin. VipA increases the polymerization of actin filaments *in vitro* by promoting the nucleation step without the requirement of additional partners, and impairs vesicle trafficking in *Saccharomyces cerevisiae*. In this work, we sought the interacting partners of VipA in early endosomes and explored the effects of this association. We found that VipA interacts directly to two key components of these organelles, the membrane lipid PI3P and the small GTPase Rab5. Binding to Rab5 requires the N-terminal region of VipA but not the C-terminal/actin-binding region. Furthermore, binding to Rab5 inhibits VipA-mediated actin polymerization by preventing *de novo* actin filament formation. These findings offer new insights into VipA’s mode of action and underscore the intricate interactions between *L. pneumophila* effectors and their host cell targets, namely the endocytic pathway.

## INTRODUCTION

*Legionella pneumophila* is an opportunistic human pathogen responsible for causing Legionnaires’ Disease, a severe pneumonia that may occur upon the inhalation of contaminated water droplets [1], [2]. The pathogen’s capacity to initiate infection in alveolar macrophages is driven by selective pressures exerted by its environmental hosts, amoebae [3]. Inside both types of professional phagocytes, *L. pneumophila* resides within a reconfigured compartment known as the *Legionella*-containing vacuole (LCV) that escapes conventional phagosomal maturation. This compartment allows access to essential nutrients, limits inter-bacterial competition and provides protection against complement-mediated lysis and antibody responses. To establish a replication-competent LCV, *L. pneumophila* inhibits phagolysosomal fusion by diverting this compartment from the endocytic pathway [4]. This requires the Icm/Dot type IV secretion system that translocates over 300 bacterial effector proteins into host cells, which control a multitude of host cell pathways by acting in a timely and spatially concerted manner [5] [6]. The repertoire of effectors presents substantial functional redundancy, meaning that multiple effectors share the same eukaryotic target and consequently are rarely essential for intracellular replication in standard experimental models [7]. Nevertheless, over the past two decades, numerous *L. pneumophila* effectors and their targets have progressively been unravelled and characterized (recently reviewed in [8], [9], [10], [11]).

Rab GTPases are key regulators of vesicle trafficking and, together with their regulators and downstream effector proteins, confer organelles their specific identity. Therefore, it is logical that Rab GTPases and related proteins are major targets of intravacuolar pathogens. Rab GTPases alternate between an active GTP-bound form and an inactive GDP-bound state and their interconversion is controlled by the action of Guanine Exchange Factors (GEFs) and GTPase-Activating Proteins (GAPs). GEFs facilitate the replacement of GDP for GTP, allowing Rabs to interact with their effectors and carry out their functions in vesicle formation, movement, and fusion. GAPs in turn increase the intrinsic GTPase activity of Rabs, promoting the conversion from the active GTP-bound state to the inactive GDP-bound state [12]. In particular, endosomal Rab GTPases are manipulated by bacteria to prevent the progressive transfer of endo-lysosomal membrane and luminal constituents to the pathogen-containing vacuoles [2]. For example, Rab5 is a key regulator of endosomal maturation by orchestrating the conversion of early endosomes into late endosomes. Active Rab5 recruits its effector VPS34, a phosphatidylinositol 3-kinase that phosphorylates phosphoinositide (PI) into phosphoinositide 3-phosphate (PI3P), contributing to forming the identity of the early endosomes and to trigger their further maturation into late endosomes [13][14][15][16].

The role of Rab5 in the development of the LCV and its modulation by *L. pneumophila* effectors has been addressed in several studies. Rab5 is localized on the surface of the LCV during early stages of host cell infection in both amoebae and macrophages [17], [18]. In addition, overexpression of Rab5 is detrimental to *L. pneumophila* replication, likely due to its impact on the integrity of the LCV membrane rather than an increase in lysosomal fusion [19] [17][20] [21]. To date, three *L. pneumophila* effectors modulating Rab5 activity have been identified. VipD has a dual Rab5 inhibitory role by preventing its recruitment to early endosomes and by blocking its access to a downstream target (early endosomal antigen 1 (EEA1), thereby contributing to avoidance of phagosomal maturation [22], [23]. Additionally, SidC/SdcA is involved in recruitment of Rab5 to the surface of the LCV and promotes its ubiquitination therein [24]. Finally, Lpg0393 is a Rab5 GEF [25]. However, the exact role of SidC/ScdA and Lpg0393 during infection remains uncertain. Thus, despite being a key player in trafficking of the LCV along the phagolysosomal pathway, the mechanisms used by *L. pneumophila* to control the action of Rab5 are mostly unknown.

Our previous studies showed that the *L. pneumophila* effector VipA (339 amino acid residues) is an actin nucleator that associates with early endosomes and actin patches during infection of macrophages, and interferes with vesicle trafficking in *S. cerevisiae* Functional analyses of truncated VipA mutants assigned a fundamental role of its COOH-terminal region (residues 206-339) in actin binding and polymerization, and the requirement of its N-terminal region (residues 1-133) for association with early endosomes. In this work, we identified PI3P and Rab5 as the endosomal anchors of VipA. Interaction of VipA with Rab5 is direct, mediated by the N-terminal region of VipA, and hinders the ability of VipA to induce *de novo* polymerization of actin filaments. These findings provide novel insights on the mode of action of VipA and highlight the complex interplay between *L. pneumophila* effectors and their host cell targets.

## MATERIAL AND METHODS

### Bacterial strains and culture media

*Escherichia coli* strains used in this work are listed in Table S1 and were routinely grown as previously described [26]. For protein overexpression and Bacterial Two-Hybrid assays see sections below.

### Plasmids and oligonucleotides

Plasmids and oligonucleotides used in this study are listed in Tables S1 and S2. For general cloning procedures, restriction enzymes, T4 DNA ligase and Phusion DNA polymerase (Thermo Fisher Scientific) were used according to the manufacturer’s instructions. The accuracy of the cloned nucleotide sequences was confirmed by DNA sequencing.

### Mammalian Cell Culture, Transfections and Infections

CHO FcγRII cells [27] were grown in Dulbecco’s Modified Eagle Medium (DMEM; Corning) and 10% (v/v) heat-inactivated fetal bovine serum (FBS; Thermo Fisher Scientific), at 37°C in a 5% (v/v) CO2 incubator. CHO cells were seeded on 24-well plates at 1×10^5^ cells/well or 5×10^4^ cells/well for, respectively, immunoblot or microscopy experiments. Cells were transfected using the jetPEI™ reagent (Polyplus) according to manufacturer’s protocol for 24 hours.

### Immunofluorescence Microscopy

Transfected CHO cells cells were fixed for 20 minutes in 4% PFA (v/v, in PBS) and permeabilized with Triton X 0.1% (v/v, in PBS) for the same time. For labelings, primary antibody (mouse anti-myc, Clone 9E10, Calbiochem), was diluted 1:200 in PBS with 10% horse serum and incubated for 1 hour, followed by PBS washing and incubation with appropriate fluorophore-conjugated anti-mouse antibodies (anti-mouse-AlexaFluor350 or anti-mouse-AlexaFluor560, Jackson ImmunoResearch) in the same conditions. DNA labeling was carried out with 4,6-Diamidino-2-phenylindole (DAPI; 1:30.000), and actin staining was carried out by incubation with Phalloidin-Alexa555 (Thermo Fisher Scientific, 1:200) during 30 min. Images were acquired on an Axio Imager D2 (Zeiss) or a Laser Scanner Confocal Microscope Zeiss LSM710 (for colocalization experiments) and processed with ImageJ 2.14 software. Quantitative analysis of co-localization between VipA and FYVE-GFP was performed by calculating the Manders overlap coefficient, corresponding to the fraction of red pixels (VipA signal) that overlaps with green pixels (FYVE-GFP) in relation to the total red pixels. For this purpose, signal intensities for each cell (n≥12, two independent experiments) were processed in ImageJ and the coefficients determined with the plugin JACoP [28]. Statistical analysis was performed using GraphPad Prism software version 5.02. P values were calculated using a t test (unpaired, two tails).

### Co-Immunoprecipitation Assays

CHO cells were transfected with pEGFP-N1, plasmids encoding EGFP-Rab5 variants, pEF6a and plasmids encoding VipA-myc variants using the jetPEI^TM^ transfection reagent (Polyplus) according to manufacturer’s protocol. Cells were harvested 24 hours post transfection and co-immunoprecipitations performed using Chromotek GFP-Trap agarose beads essentially as described by the manufacturer. In brief, cells were lysed with lysis Buffer (50 mM Tris-HCl pH 8, 150 mM NaCl, 0.1 mM EDTA, 0.5% [w/v] IGEPALÒ CA-630, 1 mM dithiothreitol [DTT], 100 μg/ml phenylmethylsulfonyl fluoride [PMSF] and a protease Inhibitor cocktail [Amresco]), and the beads were incubated with lysate-supernatants for 4 hours at 4 °C in dilution buffer (50 mM Tris-HCl pH 8, 150 mM NaCl, 0.1 mM ethylenediaminetetraacetic acid [EDTA], 1 mM DTT, 100 μg/ml PMSF and a protease Inhibitor cocktail [Amresco]), and washed three times with wash buffer (50 mM Tris-Hcl pH 8, 150 mM NaCl, 0.1 mM EDTA). Aliquots of input, non-bound, washes and bound samples were analyzed by immunoblotting (see below).

### SDS-PAGE and immunoblotting

For SDS-PAGE, samples containing 1x Laemmli buffer were boiled for 5 minutes and ran on 12% gels at 150 V for 75 minutes. Gels were processed for immunoblotting using Trans-Blot Turbo Transfer System (BioRad) and 0.2 μm pore-size nitrocellulose membranes (BioRad). Membranes were incubated 1 hour with primary antibodies diluted 1:1000 in Blocking solution (PBS with 0.1% Tween and 4% non-fat milk powder; goat anti-GFP, SICGEN; rabbit anti-VipA; mouse anti-myc, Clone 9E10, Calbiochem), followed by PBS washing and incubation with appropriate secondary antibodies in the same conditions (anti-goat or anti-mouse horseradish peroxidase (HRP)-conjugated secondary antibodies, GE Healthcare or Jackson ImmunoResearch; 1:10000). For VipA-lipid interaction, PIP strips membranes (Echelon Biosciences) were incubated with 5 mg of purified His_6_-VipA in Blocking solution and hybridation carried out as above. Immunoblot detection was done with Western Lightning ECL Pro (Perkin Elmer) and exposure to Amersham Hyperfilm ECL (GE Healthcare).

### Overexpression and purification of His_6_-VipA and GST-Rab5 proteins

*E. coli* BL21(DE3) strains harboring plasmids encoding 6xHis-tagged proteins (see Table S1) were grown at 37°C for 18 hours (His_6_-VipA and His_6_-VipA_ΔNH2_) or at 37°C for 5 hours followed by 24 hours at 26°C (His_6_-VipA_ΔCOOH_) in auto-induction conditions [29]. Cells were harvested by centrifugation and the cell pellet was resuspended in 10 ml of lysis buffer (50 mM Na_2_HPO_4_, 300 mM NaCl, 20 mM imidazole). Bacteria were lysed with 3 passages in a French press at 900 PSI in the presence of 1 mM PMSF and lysates were centrifuged at 13000 xg for 30 minutes at 4°C. His-tagged proteins were purified from the soluble fraction using Ni-NTA (nickel-nitrilotriacetic acid) chromatography (Qiagen) and eluted with Imidazole gradients. For purification of GST-Rab5 proteins (GST-Rab5^WT^, GST-Rab5^CA^ or GST-Rab5^DN^), the corresponding strains were grown in LB and overexpression induced for 4 hours by addition of 1 mM IPTG. Bacteria were then pelleted, resuspended in PBS and lysed as above. Proteins were purified by loading the soluble fractions on glutathione-sepharose columns and elution with 10 mM glutathione in PBS. Where indicated, loading of Rab5 with with GTPgS, a non-hydrolysable form of GTP, was carried out essentially as described [30] with the following alterations. Purified GST-Rab5 was incubated for 30 min at room temperature with 2 ml of Nucleotide Exchange Buffer (20 mM HEPES pH 7.5, 100 Mm NaCl, 10 mM EDTA, 5 mM MgCl_2_, 1 mM DTT) containing 1 mM GTPgS (Jena Bioscience) in a final 3 μM GST-Rab5 concentration. The reaction mixture was transferred to a 6 ml Centrifugal Concentrator column (MWCO 10 kDa, Vivaspin), subjected to two centrifugation cycles at 4°C and 5,000 x g for 2 min. Then, 2 ml of Nucleotide Stabilization Buffer supplemented with 10 μM of GTPgS were loaded into the column, which was centrifuged twice at 4°C, 5,000 x g for 2 min, and once at 2,000 x g for 2 min. The remaining ∼ 200 μl of protein solution was recovered and stored at −20°C until further use.

### Bacterial Two-Hybrid Assays

The bacterial adenylate cyclase two-hybrid (BACTH) system [43] (Euromedex) was used. This system includes four plasmids (pUT18, pUT18C, pKNT25 and pKT25; Table S1) enabling fusions to the N- or C-terminus of T18 or T25 fragments of the catalytic domain of adenylate cyclase (CyaA) from *Bordetella pertussis*. For bacterial two-hybrid, adenylate cyclase (Cya)-deficient *E. coli* BTH101 was transformed with plasmids expressing the different T18 and T25 hybrid proteins. Qualitative screening of putative interactions was firstly made on plates by growing co-transformants on solid LB supplemented with isopropyl β-D-1-thiogalactopyranoside (IPTG; 0.5 mM), the chromogenic substrate 5-bromo-4-chloro-indolyl-β-D-galactoside (X-gal; 40 µg/ml) and appropriate antibiotics. Plates were incubated for 24 h at 28°C. Quantification of β-gal activity was accomplished by monitoring the β-gal-dependent cleavage of *ortho*-nitrophenyl-β-galactoside (ONPG) into the colorimetric molecule *ortho*-nitrophenol, as described by the manufacturer (Euromedex). In all experiments, we used as positive control strain BTH101 transformed with plasmids pUT18C-zip and pKT25-zip, encoding two leucine zipper domains, and the indicated negative control strains. The final results were based on at least three independent experiments, using three different colonies. Statistical significance was assessed using a student t-test as indicated.

### *In vitro* actin polymerization assays

Preparation of G-Actin and polymerization assays were carried out essentially as described [31] with the following alterations. Samples contained 3 μM G-actin (Cytoskeleton, >99% pure; 20% Pyrene-actin) and purified His_6_-VipA mutants in G-Mg buffer were added as indicated. Polymerization assays were carried out in quartz cuvettes (3 mm optical path length), and fluorescence read in a Varian Cary Eclipse fluorescence reader. Values were obtained using an excitation wavelength of 365 nm and emission of 407 nm (10 nm slit width) and recorded at 1 min intervals. Data were collected with Cary Eclipse software and then processed in Excel (Microsoft).

For quantification of the number of filaments by microscopy, handmade flowcells were prepared as a sandwich of a glass slide (1.0–1.2 mm thick, Fisherbrand) with two Parafilm strips placed longitudinally over it with an approximately 15 mm wide channel between them and a glass coverslip (20 x 20 mm, Menzel-Gläser) on top. This set was heated at 100 °C to melt the parafilm. Liquid was introduced using a micropipette on one end of the channel and pulled through the flow cell by using an absorptive cloth on the other end. Before sample introduction, the flow cells were incubated with 50 μl of 10% (w/v) BSA for 10 min to reduce non-specific adsorption of actin to the surfaces of the glass. Flow cells were washed with 90 μl of G-buffer (1 mM Tris-HCl pH 7.8, 0.2 mM CaCl_2_, 0.5 mM DTT, 0.2 mM Mg-ATP) and then 20 μl of filamin (0.1 mg/ml, from turkey smooth muscle, Hypermol) was applied as tethering protein and left to incubate for 5 minutes. Finally, flow cells were washed again with 90 μl of G-buffer.

*In vitro* actin polymerization reactions were performed as follows: 100 nM Atto488-Actin (α-skeletal muscle actin from rabbit skeletal muscle, Hypermol) was incubated for 5 minutes on ice with G-buffer and 5% (v/v) of 10x ME buffer (4 mM MgCl_2_, 20 mM EGTA) to exchange Ca^2+^ and Mg^2+^. His_6_-VipA, GST-Rab5 and G-buffer were then added to a final volume of 100 μl. Reactions were initiated by introducing 10x KMEI (500 mM KCl, 20 mM MgCl_2_, 20 mM EGTA, 300 mM imidazole) and 40% (v/v) of Imaging Buffer (40 mg/ml catalase from bovine liver, 1.8 mg/ml glucose oxidase from *Aspergillus niger*, 100 mM a-D-glucose, 2.5% b-mercaptoethanol, 0.5% methylcellulose). After introducing the different components for each sample, both open ends of the channel were sealed with grease (korasilon-paste, GmbH).

Widefield imaging was done on a Leica DMI6000 inverted microscope, using illumination from a Leica EL6000 source, a fluorescence filter cube with the Leica GFP ET filter set, a 100x/1.46 a-plan apochromat oil immersion objective plus a 1.6x magnifier, Leica type F immersion oil, and an Evolve 512 electron microscopy charge-coupled device (EM-CCD) camera (Photometrics) using 16-bit EM gain amplification. The final pixel size was 100 nm. To assess actin polymerization over time, one-hour time-lapse videos were obtained starting five minutes after sample preparation. To evaluate nucleation at early time points with higher number of images, at least 20 images of different fields of view were acquired for every sample five minutes after sample preparation. To automatically quantify the number of actin filaments, fluorescence images were segmented using DeLTA [32], using model weights that we obtained from training the DeLTA neural network on manually segmented training data. Binary images were then processed using the Ridge Detection plug-in in ImageJ (https://imagej.net/plugins/ridge-detection) to obtain the number and lengths of filaments in individual images. This data was then analyzed and plotted using custom Python scripts.

## RESULTS

### VipA interacts with the early endosome membrane lipid PI3P

Our previous results showed a partial localization of VipA to early endosomes when ectopically expressed in mammalian epithelial cells and when delivered into THP-1 macrophages during infection by *L. pneumophila* [26], [31]. To gain insight into the function of VipA in these organelles, we were interested in finding its molecular target(s) therein. Two key identifiers of early endosomes are the lipid PI3P and the small GTPase Rab5, fundamental in their maturation process. We initially assessed a possible association with PI3P by looking at the localization of ectopically expressed VipA-myc and the PI3P probe 2xFYVE-GFP in cotransfected CHO cells, a reliable model to mimic the VipA localization observed during infection [26]. We observed a partial overlap between vesicles enriched in PI3P (PI3P^+^ vesicles) and VipA_WT_ puncta (Fig. 1A), similar to the partial colocalization between vesicles enriched in early endosome marker EEA1 (Early Endosome Antigen 1) and VipA we had found previously [31].

**Figure 1.**
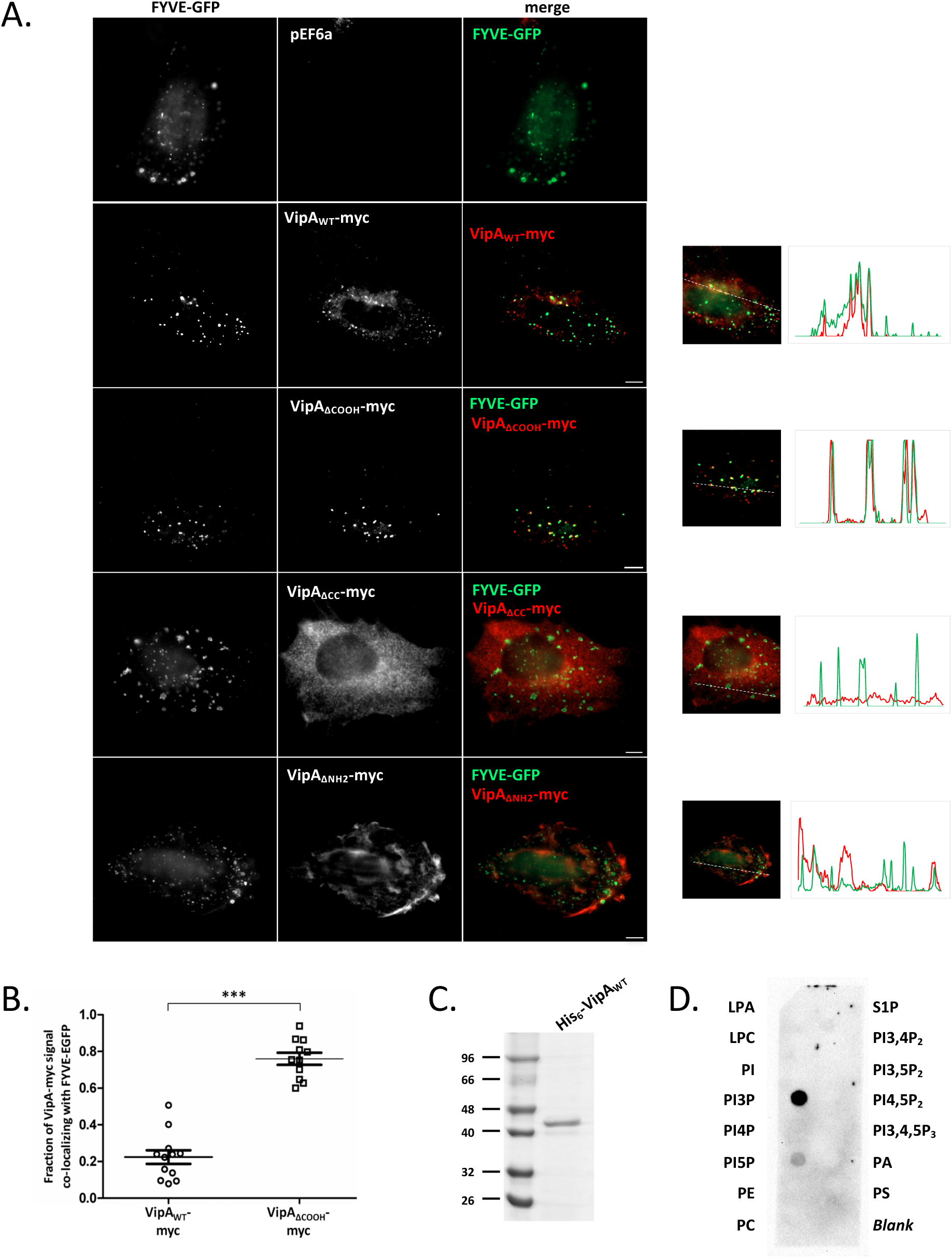
*L. pneumophila* effector VipA interacts with lipid PI3P in CHO cells and *in vitro*. **A.** CHO cells were cotransfected with plasmid pEF6a derivatives encoding myc-tagged versions of VipA (wild-type, VipA_WT_; lacking aminoacid residues 207-339, VipA_ΔCOOH_; lacking aminoacid residues 134-206, VipA_ΔCC_; or lacking aminoacid residues 1-133, VipA_ΔNH2_; in red), and plasmid pEGFP-N1 derivatives encoding GFP or FYVE-GFP (in green). Cells were fixed with 4% PFA, permeabilized with 0.1% Triton-X and labeled with an anti-myc antibody (mouse) and secondary anti-mouse-Rhodamine. Microscopy was performed on a Laser Scanner Confocal Microscope (Zeiss LSM710). Representative images are shown (left) and corresponding Profile Plots for the intensity of FYVE-GFP and VipA were performed with ImageJ (right). Scale bars, 10 µm. **B.** Quantification of colocalization was performed for VipA_WT_ and VipA_ΔCOOH_ by determining the fraction of the effector overlapping with FYVE-GFP in each cell (Manders coefficient). Bars correspond to mean and standard error of the mean; _***_, P<0.001. **C.** Coomassie stained SDS-PAGE with 2.2 ug of Ni-NTA purified His_6_-VipA_WT_. **D.** Membrane containing the indicated immobilized lipids was incubated with purified His_6_-VipA and probed with polyclonal anti-VipA (rabbit) and secondary anti-rabbit-HRP (goat) antibodies. LPA, Lysophosphatidic Acid; LPC, Lysophosphocholine; PI, PhosphatidyIinositol; PI3P, PhosphatidyIinositol 3-Phosphate; PI4P, PhosphatidyIinositol 4-Phosphate; PI5P, PhosphatidyIinositol 5-Phosphate; PE, Phosphatidylethanolamine; PC, Phosphatidylcholine; S1P, Sphingosine-1-phosphate; PI3,4P_2_, PhosphatidyIinositol 3,4-Bisphosphate; PI3,5P_2_, PhosphatidyIinositol 3,5-Bisphosphate; PI4,5P_2_, PhosphatidyIinositol 4,5-Bisphosphate; PI3,4,5P_3_, PhosphatidyIinositol 3,4,5-trisphosphate; PA, Phosphatidic Acid; PS, Phosphatidylserine.

To find the region of VipA involved in this process, we used VipA-myc constructs previously characterized [31], either lacking the N-terminal region (mutant VipA_ΔNH2_, deletion of amino acid residues 1-133), the central coiled-coil region (VipA_ΔCC_; lacking a.a. 134-205), or the C-terminal actin-binding region (VipA_ΔCOOH_; missing a.a. 206-339). Removal of the two N-terminal and coiled-coil regions resulted in no obvious colocalization with FYVE-GFP and caused to a loss of the typical VipA puncta, leading a mostly cortical localization for VipA_ΔNH2_ and cytosolic distribution for VipA_ΔCC_ (Fig. 1A). In contrast, and strikingly, deletion of the actin-binding region of VipA led to its conspicuous association with PI3P^+^ vesicles, reflected in the concomitant red and green peaks in the intensity profile plot for both signals (Fig. 1A). In the case of the two VipA variants displaying signal overlap with PI3P, VipA_WT_ and VipA_ΔCOOH_, the degree of colocalization was quantified by determining the fraction of VipA overlappping with FYVE-GFP in a cell (Manders coefficient). Confirming the previous visual examinations, the average degree of colocalization of VipA_ΔCOOH_ with PI3P was 76%, significantly higher than the 22% found for VipA_WT_ (Fig. 1B).

To confirm specific binding of VipA to PI3P we performed *in vitro* lipid-protein interaction assays. To do this, we incubated a membrane spotted with different phosphoinositide lipids with purified histidine-tagged VipA (His_6_-VipA_WT_; Fig. 1B). Subsequent detection with anti-VipA antibodies revealed the direct interaction of the effector with PI3P, and residually with PI5P, but not with the remaining membrane lipids (Fig. 1C). Together, colocalization of VipA with PI3P^+^ vesicles in CHO cells and direct binding to PI3P *in vitro* indicate that VipA interacts specifically with endosomal lipid PI3P.

### VipA interacts with GTPase Rab5

A subset of early endosomal proteins that bind directly PI3P, including EEA1, also interact with Rab5. Thus, we decided to test the association of VipA with this GTPase. As above, CHO cells were co-transfected with plasmids encoding VipA_WT_-myc, and expressing GFP or GFP-tagged Rab5a variants. As previously reported, in cells overexpressing GFP-Rab5 we observed the formation of enlarged Rab5^+^ endosomes due to excessive homotypic fusion of early endosomes (Fig. 2A) [33]. In addition, in these cells there was a change in the localization of VipA to the larger endosomes enriched in Rab5, instead of the small VipA puncta seen in cell producing GFP alone (Fig. 2A). To examine a possible preference of VipA to the active/GTP-bound or to the inactive/GDP-bound form of Rab5, we expressed two well-characterized GTPase mutants that mimic these two activation states. Mutant Rab5^Q79L^ is locked in its GTP-bound conformation and is therefore <u>c</u>onstitutively <u>a</u>ctive (henceforth named Rab5^CA^) whereas Rab5^S34N^ is an inactive <u>d</u>ominant-<u>n</u>egative mutant (Rab5^DN^) [15], [34]. When Rab5^CA^ was present, as observed above for cells overexpressing GFP-Rab5, VipA changed its localization from small widespread puncta to the typical enlarged endosomes resulting from the activity of this active GTPase (Fig. 2A). These endosomes are enriched in actin filaments, independently of the presence of VipA (insets for comparison in Fig. 2A). In contrast, inactive Rab5^DN^ is mostly dispersed in the cytosol with some enrichment in the perinuclear region. In cells expressing Rab5^DN^, the striking redistribution of VipA mirroring the localization of the Rab5^DN^ was not observed, although in some cells accumulation of VipA puncta in the perinuclear zone was apparent (Fig. 2A).

**Figure 2.**
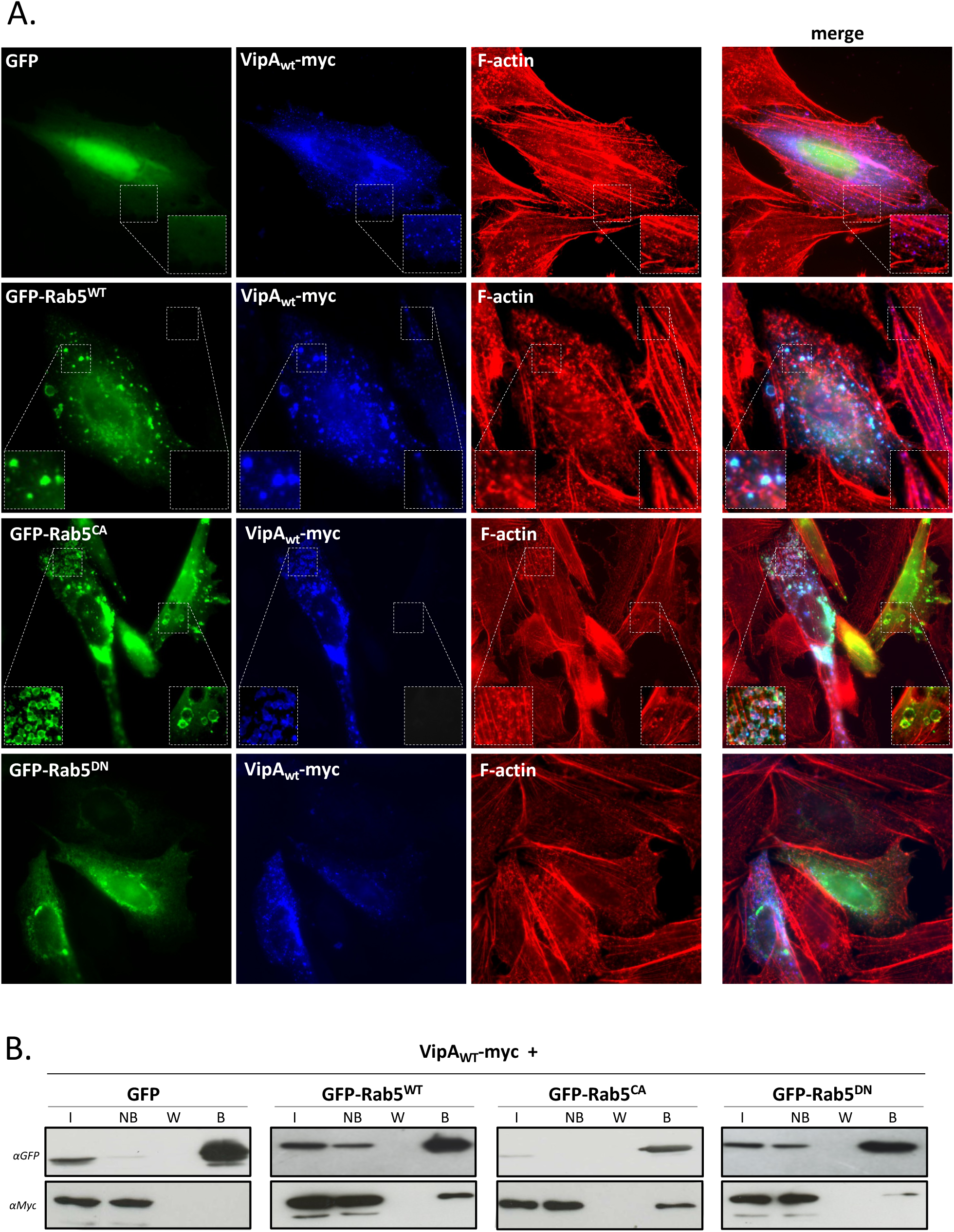
*L. pneumophila* VipA interacts with Rab5 in CHO cells. Cells were cotransfected with plasmids encoding full-length VipA-myc and wildtype or Rab5 mutants (Rab5_WT_, Rab5_CA_ or Rab5_DN_; GFP tagged, in green). **A.** Cells were fixed with 4% PFA and permeabilized with 0.1% Triton-X; VipA was labeled with an anti-myc antibody (mouse) and secondary anti-mouse-Alexa Fluor 350 (in blue), and F-actin with Phalloidin-Alexa Fluor 555 (red). **B.** For co-immunoprecipitation assays, cells were lysed and used in GFP-trap assays. Samples of Input (I), Non-bound (NB), final Wash (W) and Bound (B) fractions were separated by SDS-PAGE. Immunoblots were carried out using anti-GFP (goat) or anti-myc (mouse) antibodies followed by corresponding secondary HRP-labeled secondary antibodies.

A possible interaction between VipA and Rab5 suggested by their subcellular co-localization was tested by co-immunoprecipitation assays. For this, we used the previous experimental setup (i.e. CHO cells co-expressing VipA_WT_-myc and GFP-Rab5 variants) and immunoprecipitated GFP tagged proteins from cell lysates using GFP-Trap beads (Chromotek). After separation by SDS-PAGE, samples were subjected to immunoblots to assess VipA co-immunoprecipitation with Rab5. VipA was detected in fractions containing GFP-Rab5^WT^, GFP-Rab5^CA^ and, to a lesser extent, GFP-Rab5^DN^, but not in a sample with GFP alone (Fig. 2B).

Taken together, results from colocalization experiments and co-immunoprecipitation assays show that ectopically expressed VipA forms a complex with the small endosomal GTPase Rab5 in mammalian cells.

### Interaction of VipA with Rab5 in vivo is direct and is mediated by the N-terminal region

To identify the region of VipA necessary for its association to Rab5, CHO cells were co-transfected with the constructs producing VipA_ΔCOOH_-myc, VipA_ΔCC_-myc and VipA_ΔNH2_-myc and with the GFP-tagged Rab5a variants. VipA_ΔCOOH_ showed a similar localization to VipA _WT_-myc in cells producing GFP alone (Fig. 3A top panel), and when co-expressed with the Rab5 versions (Rab5^WT^, Rab5^CA^, Rab5^DN^), VipA_ΔCOOH_ also behaved similarly to VipA_WT_-myc, as its localization was altered from small puncta to larger organelles, distinctively overlapping with Rab5^WT^ and Rab5^CA^ (Fig. 3A, top panels). In agreement, co-immunoprecipitation assays confirmed the ability of VipA_ΔCOOH_ to form a complex with Rab5 (Fig. 3B, top panels). In the case of VipA_ΔCC_, some localization in endosomes enriched in Rab5 was visualized, as well as a residual complex formation with Rab5^WT^ (Figs. 3A and 3B, middle panels). In contrast, deletion of the N-terminal of VipA abrogated this association, as VipA_ΔNH2_ is absent in Rab5 immunoprecipitated fractions and its subcellular localization remains mostly cytosolic in the presence of the Rab5 variants (Figs. 3A and 3B, bottom panels).

**Figure 3.**
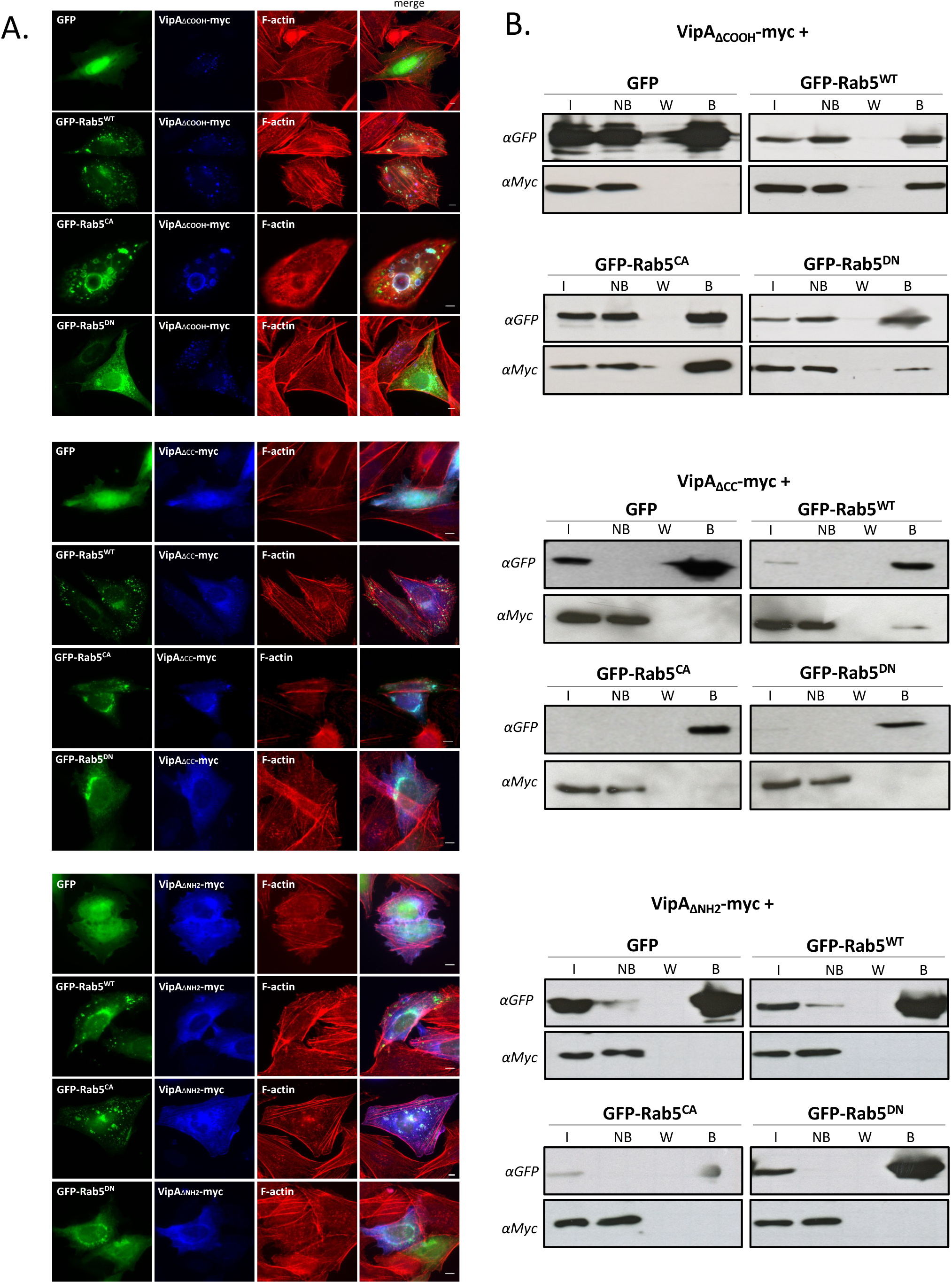
Interaction between VipA and Rab5 is mediated by the NH_2_ region of VipA. CHO cells were cotransfected with plasmids encoding VipA-myc constructs (VipA_ΔCOOH_, VipA_ΔNH2_ and VipA_ΔCC_) and wildtype or Rab5 mutants (Rab5_WT_, Rab5_CA_ or Rab5_DN_; GFP tagged). Cells were processed for microscopy analysis (**A**) or co-immunoprecipitation assays (**B**), as described in the legend of Figure 2. Scale bar, 5 μm.

To further validate the interaction of VipA with Rab5 and rule out the requirement of an additional interactant for the formation of a Rab5-VipA complex, we used a Bacterial Two-Hybrid System (BACTH; Euromedex). This methodology is based on *Bordetella pertussis* adenylate cyclase CyaA being only catalytically active when its two subunits, T18 and T25, are in physical proximity. By expressing fusion proteins containing the T18 or the T25, possible interactions will activate several reporter genes whose activity can be analyzed in specific culture media. Hence, we constructed plasmids encoding fusion proteins of wild-type VipA or Rab5 to T18 or T25 (Table S1). The reporter strain *E. coli* BTH101 was transformed with plasmid pairs expressing the two putative binding partners in all possible combinations and transformants were inoculated on LB plates containing X-gal. We observed a weak but discernible Lac^+^/blue phenotype for the strain expressing T18-VipA/T25-Rab5, suggesting an interaction between the chimeric proteins containing VipA and Rab5 at the C-terminal end (Fig. 4, left). Based on this combination of plasmids, we further investigated if complex formation was affected in variants Rab5^CA^ or Rab5^DN^. A similar weak interaction was observed with Rab5^CA^, but not with inactive variant Rab5^DN^. Additionally, and suggesting a possible inhibitory role of the actin-binding region, the interaction with Rab5^WT^ and particularly with Rab5^CA^ was significantly enhanced when variant VipA_ΔCOOH_ was used, confirmed by quantification of the β-galactosidase activity of the corresponding strains (Fig. 4, right).

**Figure 4.**
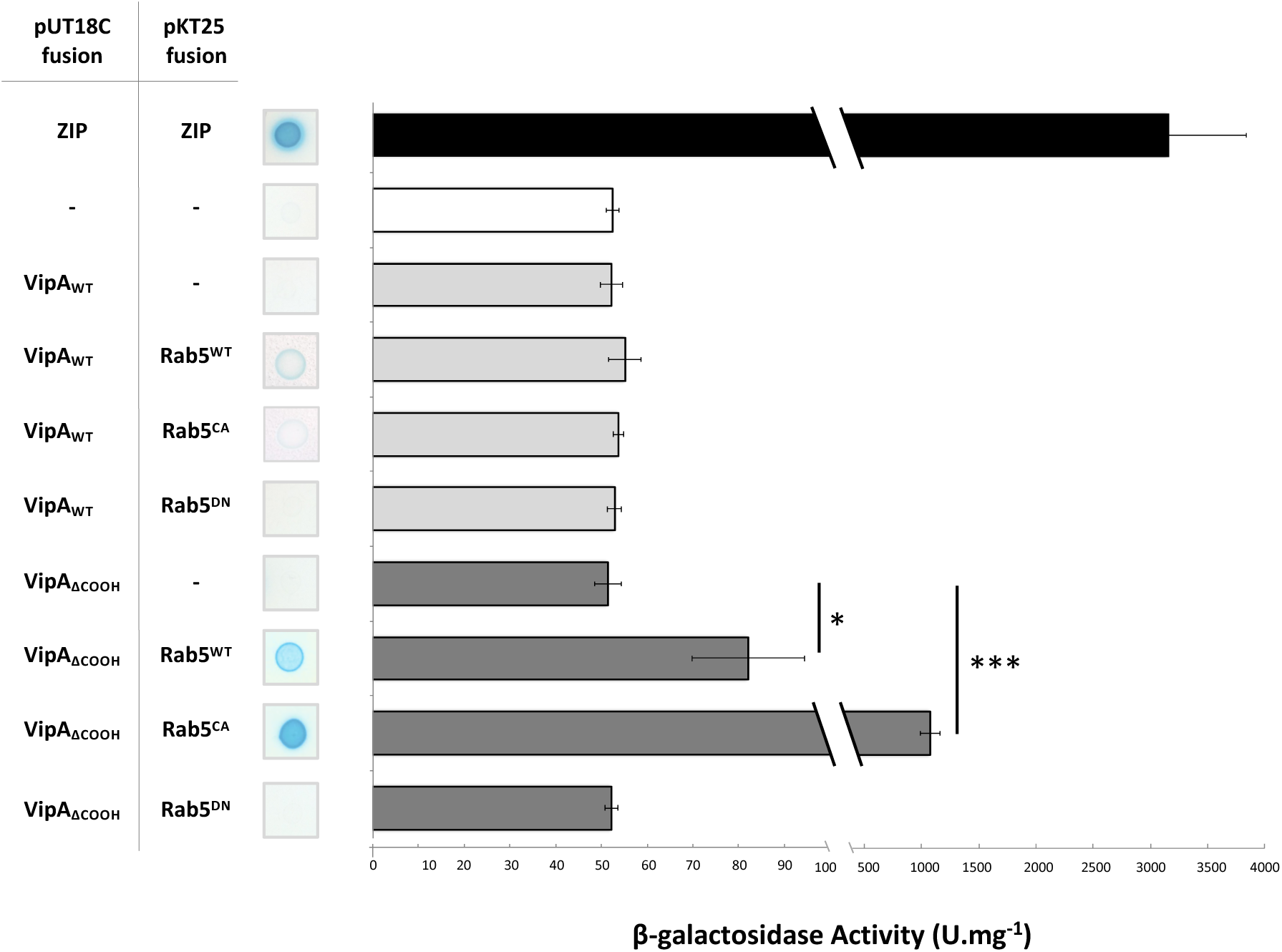
VipA interacts directly with Rab5 in a Bacterial Two-Hybrid (BACTH) Assay. *E. coli* recipient strain BTH101 was co-transformed with plasmids encoding a Rab5 variant (Rab5^WT^, Rab5^CA^ or Rab5^DN,^ encoded in plasmid pUT18C) and a VipA variant (VipA_WT_ or VipA_ΔCOOH,_ encoded in plasmid pKT25). A positive control for the BACTH system was used (Zip-Zip interaction, black bar). Negative controls included a strain with the two empty vectors (white bar) and strains expressing a protein under test and the counterpart empty vector. Activity of reporter gene *lacZ* was measured qualitatively by plating in solid media with X-gal (left) or quantitatively by β-galactosidase assays in liquid media (right) or. Bars and error bars represent the average and standard error of the media of at least 3 independent experiments. Statistical analysis was made using a Student t-test (*, p<0.05; ***, p<0.001).

Taken together, the results from microscopy, co-immunoprecipitation and bacterial two-hybrid assays indicate that VipA and Rab5 form a complex in mammalian cells that does not require additional eukaryotic binding partners. The interaction is favored by the active form of Rab5, requires the N-terminal region of VipA but a possible inhibitory mechanism may be conveyed by the C-terminal/actin-binding region of VipA.

### Active Rab5 inhibits VipA-mediated de novo formation of actin filaments

We tested a possible effect of the binding of Rab5 to VipA on the ability of VipA to nucleate the polymerization of new actin filaments. *In vitro* pyrene-actin polymerization assays were performed as previously described [31] in the presence of purified His_6_-VipA and the three aforementioned Rab5 variants carrying an N-terminal GST tag. The presence of active mutant Rab5^CA^ and, to a lesser extent, of Rab5^WT^, negatively affected the capacity of wild-type VipA to drive F-actin polymerization (Figs. 5A and 5B), whereas the inactive GTPase Rab5^DN^ did not (Fig. 5C). This inhibitory effect of Rab5^CA^ was not observed in the case of VipA_ΔNH2_ (Fig. 5D), a mutant that still retains the ability to polymerize actin filaments. This result can be is consistent with our other results suggesting that Rab5-VipA interaction is mediated by the N-terminal region of VipA.

**Figure 5.**
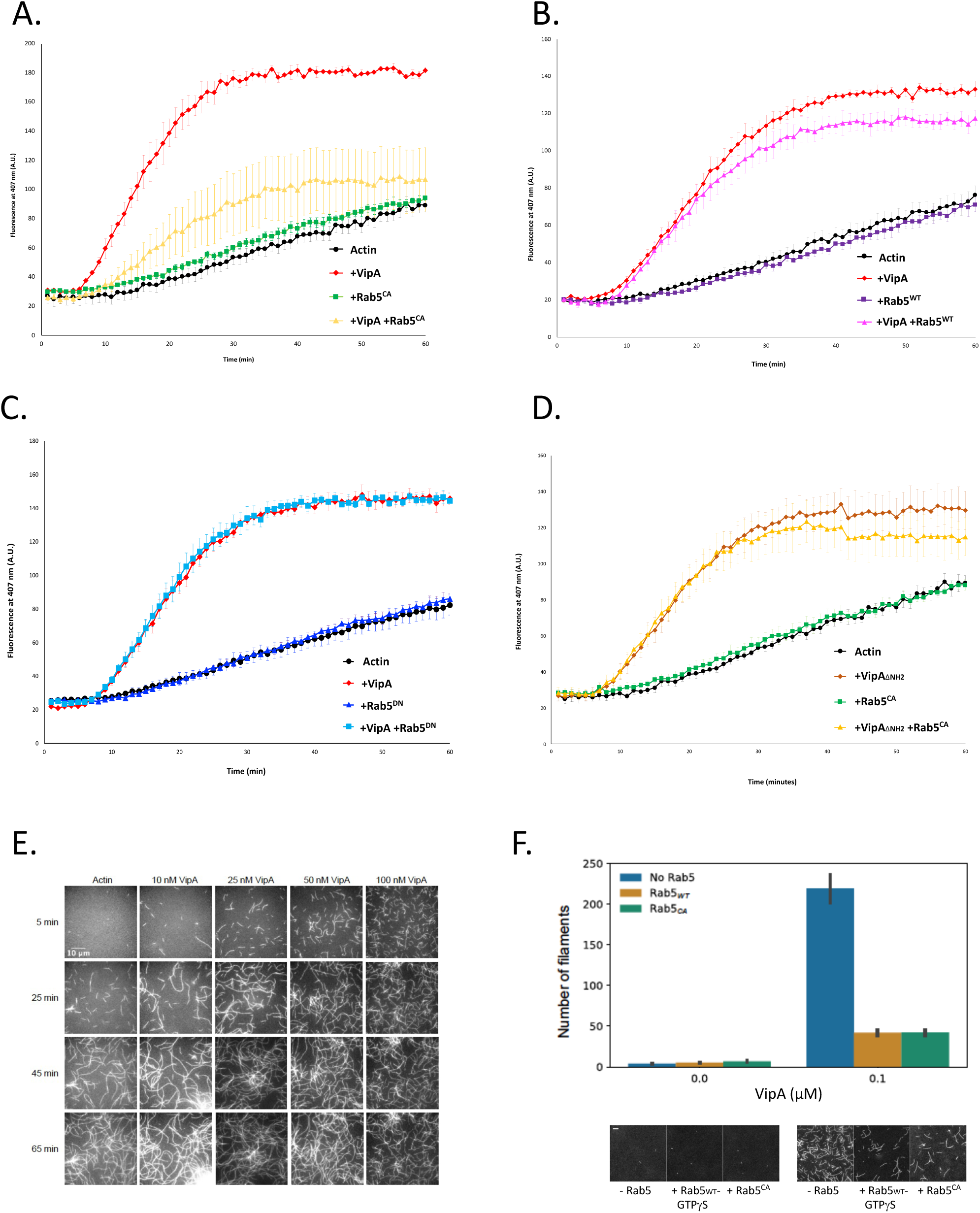
Rab5 inhibits VipA-mediated actin polymerization *in vitro*. **A-D,** In vitro pyrene-actin assays were carried out as described (details in Materials and Methods) in the presence of 3 µM of G-actin (20 % pyrene-labeled), in the presence of 500 nM of the indicated variants of purified VipA (His_6_-VipA_WT_ or His_6_-VipA_ΔNH2_) and Rab5 (GST-Rab5_CA_, GST-Rab5_WT_ or GST-Rab5_DN_). Results represent the average of three independent assays and error bars the standard error of the mean. **E,** Microscopic analysis of actin polymerization in the presence of VipA. Samples contained 100 nM of Atto 488-monomeric actin and the indicated His_6_VipA concentrations, and for each condition the same field of view was imaged at 5, 25, 45 or 65 minutes after reaction start (representative images are shown). **F**, Quantitative analysis of the number of actin filaments at 5 minutes after reaction start, in the absence or presence of 100 nM of VipA, Rab5_WT_ or Rab5_CA_. Results represent the average number of filaments counted in at least 20 fields of view from 3 independent experiments, bars correspond to a 68% confidence interval. Representative images of each condition shown below the corresponding quantification; scale bar, 10 μm.

To confirm that this bulk effect on actin polymerization was due to a decrease in VipA-mediated actin nucleation, we quantified *de novo* actin filament formation. This was carried out by microscopic analysis and counting the number of filaments polymerized in reactions containing Atto 488-monomeric actin and different concentrations of VipA and Rab5 at different timepoints. VipA increased the number of actin filaments in a concentration (0-100 nM) and time-dependent (0-65 min) manner (Fig. 5E), in agreement with its actin nucleator function. To assess the effect of Rab5 on the capacity of VipA to nucleate F-actin *de novo*, we quantified the number of existing filaments at the earliest measurable time-point, 5 minutes. In these conditions, a 50-fold increase in the amount of filaments is observed when 100 nM of VipA is present (Fig. 5E, quantification in Fig. 5F). This reaction was then repeated in the presence of purified GST-Rab5^CA^ or GST-Rab5^WT^ loaded with the GTP analog GTPγS, that locks the GTPase in the active conformation. These Rab5 GTPases did not affect *de novo* F-actin polymerization, but both significantly reduced by 5-fold the ability of VipA to nucleate new filaments (Fig. 5F).

Overall, *in vitro* actin bulk polymerization assays together with qunatification of the number of actin filaments show that binding of active Rab5 to VipA strongly reduces this effector’s ability to nucleate *de novo* F-actin polymerization.

## DISCUSSION

In this study, we advanced the understanding of the *L. pneumophila* effector VipA by identifying novel binding partners in eukaryotic cells. Beyond its previously established interaction with actin, we showed that VipA binds to the phospholipid PI3P and the small GTPase Rab5, two critical components of early endosomes. The interaction between VipA and Rab5 is direct and mediated by the N-terminal region of VipA. Furthermore, we observed that binding to the active form of Rab5 inhibits VipA-mediated *de novo* actin filament formation, unveiling a downstream effect of the VipA-Rab5 interaction.

Rab GTPases, including Rab5, have emerged as key targets of intracellular bacteria. In *L. pneumophila*, a deleterious effect of Rab5 in bacterial replication has been highlighted by gene silencing studies in A549 epithelial lung cells and in bone marrow-derived macrophages [17], [19]. Reports on the presence of Rab5 on the LCV have been divergent. Initial studies with HeLa cells overexpressing Rab5c suggested that phagosomes containing wild-type *L. pneumophila* did not acquire this GTPase [35]. However, later proteomic analyses of the LCV membrane revealed the presence of all Rab5 isoforms in the LCVs of macrophages and model amoeba *Dictyostelium discoideum*, a localization that is at least partially dependent on a functional Icm/Dot T4SS [17] [18].

Previous studies have identified two Icm/Dot effectors as Rab5 regulators. Translocated VipD binds to the GTP-bound active form of Rab5 on early endosomes in the vicinity of the LCV, blocking its interaction with downstream effectors. Together with VipD’s phospholipase A1 activity, this interaction disrupts endosomal trafficking, thereby reducing LCV exposure to the endosomal compartment and ultimately preventing *Legionella*’s lysosomal degradation in macrophage cells [22], [23]. Another effector, Lpg0393, functions as guanine nucleotide exchange factor (GEF) for endosomal Rab5, Rab21 and Rab22, although the physiological relevance of this activity during infection remains unclear [25]. Interestingly, in the *L. pneumophila* strain Philadelphia-1 genome, *lpg0393* is located near *vipA* (*lpg0390*), suggesting a gene cluster encoding effectors involved in modulation of the Rab5 pathway.

Recent insights into VipA’s function have been expanded by a large-scale Yeast Two-Hybrid screen, which identified two potential novel VipA binding partners [36]. The first is the *L. pneumophila* metaeffector Lpg2149, which blocks the activity of effectors MavC and MvcA involved in β-actin ubiquitination and NFkB activity [36] [37]. The second is the eukaryotic protein CDK4, a serine/threonine kinase crucial for cell cycle progression [38]. Further studies have revealed LegK2-VipA as an effector-effector suppression pair [39]. Although these effectors do not physically interact, deletion of *vipA* rescued the defects in intracellular replication and endosomal escape observed in *ΔlegK2* mutants in *A. castellanii*. Interestingly, a role for VipA during the preceding bacterial internalization step was also observed. In fact, in lung epithelial cells, VipA promoted the elongation of membrane wraps around filamentous *L. pneumophila* triggered by bacterial adhesion to host cell β1 integrin and E-cadherin receptors facilitating entry [40].

Collectively, these findings reveal a complex network of VipA interactions with both bacterial and eukaryotic proteins, underscoring its importance in *L. pneumophila’s* the interaction with host cells. Further studies are needed to elucidate the molecular mechanisms used which VipA modulates actin dynamics and key endocytic components to coordinate, alongside other effectors, the manipulation of endolysosomal maturation.

## ACKNOWLEDGMENTS

This project has been funded by: Fundação para a Ciência e a Tecnologia, I.P. (FCT) Grant PTDC/BIA-MIC/2821/2012 to I.F, FCT National funds in the scope of the project UIDP/04378/2020 and UIDB/04378/2020 of the Research Unit on Applied Molecular Biosciences-UCIBIO and the project LA/P/0140/2020 of the Associate Laboratory Institute for Health and Bioeconomy - i4HB. Z.H. is supported by Fundação para a Ciência e a Tecnologia (FCT) through MOSTMICRO-ITQB (DOI 10.54499/UIDB/04612/2020; DOI 10.54499/UIDP/04612/2020) and LS4FUTURE Associated Laboratory (DOI 10.54499/LA/P/0087/2020). We thank Joana Almeida for the construction of plasmid pJA6.

