## Supplemental Data for "The small GTPase Rab5 inhibits actin polymerization mediated by the *Legionella pneumophila* effector VipA"

**TABLE S1.** Bacterial strains and plasmids used in this work.

| *E. coli* strains | Purpose | Reference |
|---|---|---|
| **NEB 10b** | Plasmid cloning | New England Biolabs |
| **BL21(DE3)** | Overexpression and purification of $His_6$- and GST-tagged proteins | [41] |
| **BTH101** | Bacterial Two-Hybrid assays | [42] |
| **Plasmid** | **Purpose/Construction** | **Reference** |
| | ***Transfection of mammalian cells*** | |
| **pEF6/Myc-His a** | Vector for C-terminal myc fusions | Invitrogen |
| **pIF328** | pEF6-*vipA*; encodes $VipA_{WT}$-myc | [31] |
| **pIF344** | pEF6-$vipA_{\Delta NH2}$; encodes $VipA_{\Delta NH2}$-myc | [31] |
| **pIF361** | pEF6-$vipA_{\Delta COOH}$; encodes $VipA_{\Delta COOH}$-myc | [31] |
| **pIF368** | pEF6-$vipA_{\Delta CC}$; encodes $VipA_{\Delta CC}$-myc | [31] |
| **pEGFP-c1** | Vector for N-terminal EGFP fusions | Clontech |
| **pIF398** | pEGFP-C1-$rab5^{WT}$; encodes EGFP-$Rab5^{WT}$ (region amplified by PCR from pmRFP-Rab5 with oligos 2072 and 2073, digested with HindIII-BamHI and inserted into pEGFP-c1 HindIII-BamHI) | This work |
| **pEGFP-C1-$rab5^{Q79L}$** | pEGFP-C1-$rab5^{CA}$; encodes EGFP-$Rab5^{CA}$ | [43] |
| **pEGFP-C1-$rab5^{S34N}$** | pEGFP-C1-$rab5^{DN}$; encodes EGFP-$Rab5^{DN}$ | [43] |
| **pmRFP-Rab5** | DNA template for $rab5^{WT}$ PCRs | [44] |
| | ***Bacterial Two-Hybrid (BACTH)*** | |
| **pUT18C** | Vector for fusions to the C-terminus of fragment T18 | [42] |
| **pKT25** | Vector for fusions to the C-terminus of fragment T25 | [42] |
| **pUT18** | Vector for fusions to the N-terminus of fragment T18 | [42] |
| **pKNT25** | Vector for fusions to the N-terminus of fragment T25 | [42] |
| **pUT18C-zip** | Positive control for BACTH assays; encodes T18-ZIP | [42] |
| **pKT25-zip** | Positive control for BACTH assays; encodes T25-ZIP | [42] |
| **pJA1** | pKNT25-rab5WT; encodes $Rab5^{WT}$-T25 (region amplified by PCR from pIF398 by PCR with oligos 2269 and 2270, digested with PstI-KpnI and inserted into pKNT25 PstI-KpnI) | This work |
| **pJA2** | pUT18-vipA; encodes VipA-T18 (region amplified by PCR from *L. pneumophila* JR32 with oligos 2266 and 2267, digested with PstI-KpnI and inserted into pUT18 PstI-KpnI) | This work |
| **pJA3** | pUT18-rab5WT; encodes $Rab5^{WT}$-T18 (region amplified by PCR from pIF398 by PCR with oligos 2269 and 2270, digested with PstI-KpnI and inserted into pUT18 PstI-KpnI) | This work |
| **pJA4** | pKNT25-vipA; encodes VipA-T25 (region amplified by PCR from *L. pneumophila* JR32 with oligos 2266 and 2267, digested with PstI-KpnI and inserted into pKNT25 PstI-KpnI) | This work |
| **pJA5** | pKT25-vipA; encodes T25-VipA (region amplified by PCR from *L. pneumophila* JR32 with oligos 2265 and 2267, digested with PstI-KpnI and inserted into pKT25 PstI-KpnI) | This work |
| **pJA6** | pKT25-$rab5^{WT}$; encodes T25-$Rab5^{WT}$ (region amplified by PCR from pIF398 by PCR with oligos 2268 and 2270, digested with PstI-KpnI and inserted into pKT25 PstI-KpnI) | This work |
| **pIF407** | pUT18C-rab5WT; encodes $Rab5^{WT}$-T18 (region amplified by PCR from pIF398 by PCR with oligos 2269 and 2270, digested with PstI-KpnI and inserted into pUT18C PstI-KpnI) | This work |
| **pIF408** | pUT18C-$vipA_{WT}$; encodes T18-$VipA_{WT}$ (region amplified by PCR from *L. pneumophila* JR32 with oligos 2266 and 2267, digested with PstI-KpnI and inserted into pUT18C PstI-KpnI) | This work |
| **pIF409** | pKT25-$rab5^{CA}$; encodes T25-$Rab5^{CA}$ (region amplified by PCR from pEGFP-C1-$rab5^{Q79L}$ by PCR with oligos 2268 and 2270, digested with PstI-KpnI and inserted into pKT25 PstI-KpnI) | This work |
| **pIF410** | pKT25-$rab5^{DN}$; encodes T25-$Rab5^{DN}$ (region amplified by PCR from pEGFP-C1-$rab5^{S34N}$ by PCR with oligos 2268 and 2270, digested with PstI-KpnI and inserted into pKT25 PstI-KpnI) | This work |
| **pIF413** | pUT18C-$vipA_{\Delta COOH}$; encodes T18-$VipA_{\Delta COOH}$ (region amplified by PCR from pIF408 with oligos 2266 and 2329, digested with PstI-KpnI and inserted into pUT18C PstI-KpnI) | This work |
| | ***Protein Overexpression and Purification*** | |
| **pGEX-4T-2** | Vector for N-terminal GST fusions | GE Healthcare |
| **pIF415** | pGEX-$rab5^{WT}$; encodes GST-$Rab5^{WT}$ (region amplified from pIF398 by PCR with oligos 2331 and 2332, digested with EcoRI-XhoI and inserted into pGEX-4T-2 EcoRI-XhoI) | This work |
| **pIF416** | pGEX-$rab5^{CA}$; encodes GST-$Rab5^{CA}$ (region amplified from pEGFP-C1-$rab5^{Q79L}$ by PCR with oligos 2331 and 2332, digested with EcoRI-XhoI and inserted into pGEX-4T-2 EcoRI-XhoI) | This work |
| **pIF417** | pGEX-$rab5^{DN}$; encodes GST-$Rab5^{DN}$ (region amplified from pEGFP-C1-$rab5^{S34N}$ by PCR with oligos 2331 and 2332, digested with EcoRI-XhoI and inserted into pGEX-4T-2 EcoRI-XhoI) | This work |

| | | |
|---|---|---|
| **pET15b-*vipA*** | pET15b-His$_6$-VipA$_{WT}$ | [26] |
| **pIF358** | His$_6$-VipA$_{\Delta NH2}$ (region amplified by PCR with oligos 1592 and 1591, digested with NdeI-BamHI and inserted into pET28b) | [31] |
| **pIF373** | His$_6$-VipA$_{\Delta COOH}$ (region amplified by PCR with oligos 1590 and 1299, digested with NdeI-BamHI and inserted into pET28b) | [31] |
| **pIF374** | His$_6$-VipA$_{\Delta CC}$ (region amplified by PCR with oligos 1590 and 1591, digested with NdeI-BamHI and inserted into pET28b) | [31] |
| **pJB9** | His$_6$-VipA$_{COOH}$ (region amplified by PCR with oligos 1775 and 1591, digested with NdeI-BamHI and inserted into pET28b) | [31] |

**Table S2.** Oligonucleotides used in this work.

| Oligonucleotide | Sequence (5'→3') [a] |
|---|---|
| 2265 | AAAACTGCAGGGATGCCTATCAGTAATGCCTTTC |
| 2266 | AAAACTGCAGGATGCCTATCAGTAATGCCTTTC |
| 2267 | AAAAGGTACCCGGAGATTTTTTTTTTCGACGGG |
| 2268 | AAAACTGCAGGGATGGCTAGTCGAGGCGCAAC |
| 2269 | AAAACTGCAGGATGGCTAGTCGAGGCGCAAC |
| 2270 | AAAAGGTACCCGGTTACTACAACACTGATTCC |
| 2328 | AAAACTGCAGGTTTATTTTAGCTCAAAAGGC |
| 2329 | AAAAGGTACCCTATGTGATTTCGCTTAGAGTTTG |
| 2330 | AAAAGGTACCCTACGCTTGTTGCGTCATGACCAG |
| 2072 | AAAAAAGCTTCTATGGCTAGTCGAGGCGCAACAAG |
| 2073 | TCCGGTGGATCCTTAGTTACTACAACACTG |
| 2331 | AAAAGAATTCCCATGGCTAGTCGAGGCGCAACAAG |
| 2332 | AAAACTCGAGTTAGTTACTACAACACTGATTCC |

[a] Restriction sites are underlined.
